# Dynamics of TCR*β* repertoires from serial sampling of healthy individuals

**DOI:** 10.1101/2022.05.11.491566

**Authors:** Iñigo Ayestaran, Jamie R. Blundell

## Abstract

T-cell receptor (TCR) repertoires provide a historical record of antigen exposure. However, the dynamics of TCR repertoires in healthy individuals remain largely uncharacterised. How much of the repertoire is under immune selection in healthy individuals? Do groups of sequences under immune selection share similar dynamics due to convergent specificity? What is the relationship between dynamic similarity and sequence similarity of TCRs? Here we develop a statistical framework for identifying clonotypes under immune selection in time series repertoire data. Applying this framework to serially sampled repertoires collected over the course of a year from 3 healthy volunteers, we are able to detect hundreds of TCRs undergoing strong immune selection whereby clonotype frequencies can change by orders of magnitude over timescales as short as a month. Clonotypes under immune selection belong to a handful of distinct dynamic clusters each of which show highly coordinated temporal behaviour suggesting a common immunogenic stimulus. Whilst a subset of clonotypes within dynamic clusters show shared amino acid motif usage, most do not, suggesting the same immunogenic stimulus elicits a diverse TCR response. Conversely, shared amino acid motif usage alone identifies far fewer clonotypes under immune selection and these clonotypes do not routinely exhibit correlated temporal behaviour. These results highlight the potential of using information contained in the *dynamics* of TCR repertoires for identifying clonotypes responding to the same immunogenic stimulus in a sequence agnostic way.

## Introduction

The T-cells of the adaptive immune system recognise antigens via T-cell receptors (TCR) (*1*). The TCR is a heterodimer composed of an *α* and a *β* chain, both of which are generated following a highly variable somatic rearrangement process known as V(D)J recombination (*2*). As this recombination happens independently in each one of the ≈ 10^11^ T-cells in the human body, this generates a vast number of distinct TCRs (referred to as the TCR repertoire) with an estimated 10^6^ − 10^8^ unique sequences at the level of the *β* chain alone (*3–7*). Multiplex PCR combined with deep amplicon sequencing has been routinely used to provide quantitative surveys of the the TCR repertoire (usually at the level of the *β* chain) (*3, 8–11*). These surveys show that TCR*β* clonotypes span multiple orders of magnitude in frequency, with most clonotypes being very rare (*6, 7*). The frequency of a clonotype depends on multiple factors, including the generation probability of a given V(D)J rearrangement (*2, 12–14*), thymic selection (*15, 16*) and clonal expansion of T-cells following antigen recognition (*17*). The temporal dynamics of clonotypes are therefore shaped by antigen exposure history, including viral infections (*18*), vaccines (*19–22*) and immunotherapy (*23–25*). This raises the prospect that TCR repertoires and their dynamics may be able to be used for the purpose of disease detection (*9, 26–28*).

An important first step in exploiting TCR repertoires as a detector of disease is developing a better understanding of which TCR sequences are important in recognising and responding to which diseases. A number of strategies have been developed to identify TCRs responding to a given antigen from a static repertoire snapshot. These include approaches based on amino acid sequence similarity and motif usage (*29, 30*), identifying sets of sequences that are over-represented relative to their generation probabilities (*14, 31*), and enrichment of disease associated public TCRs in large case-control cohorts (*9, 27*). However, each of these approaches have associated limitations. Methods based on sequence similarity are limited by an incomplete understanding of the mapping between MHC-peptide complexes and the TCRs that bind them. Strategies based on identifying disease associations in large case-control cohorts are restricted to the small fraction of the sequences in the repertoire that are highly public, potentially missing signal from the large number of private clonotypes.

Longitudinal data in which repertoires are generated from the same individual over time have the potential to overcome some of these challenges by identifying clonotypes with shared dynamic behaviour (e.g. those undergoing a synchronised clonal expansion (*32*)). However, most longitudinal analyses have been focused on detecting repertoire changes in response to a specific immune stimulus (e.g. a vaccine) across small numbers of samples collected over short timescales (*18–22*). Because a common pathogen likely drives a highly coordinated temporal response in the TCRs that recognise it, analysing repertoire dynamics across longer periods of time with many samples has rich information that can be used to identify disease-related clonotypes in a sequence-agnostic way. However, to achieve this requires a three key elements. First, one requires highly quantitative diverse repertoires sampled over many time points and generated in a standard way. Second, one needs to be able to distinguish TCRs that are under immune selection from those that are fluctuating purely due to technical noise. Third, one requires robust ways of grouping sequences exhibiting highly correlated dynamic behaviour above what would be expected by chance.

Here we develop a statistical model of technical noise in TCR repertoire data and use this to identify outlier clonotypes that are under immune selection. We then develop a “dynamic similarity” metric to rationally group clonotypes into clusters with a shared temporal behaviour. This approach allows us to ask whether clonotypes within a dynamic cluster exhibit similarities at the sequence level, and conversely, whether clonotypes with shared sequence features show similar dynamics. Our study highlights the potential of quantitative longitudinal TCR repertoire analysis for the detection of broader families of public and private disease-associated TCRs and raises some potential limitations of using purely static sequence similarity based approaches when analysing full repertoires.

## Results

### Detecting clonotypes under immune selection

To examine the dynamics of TCR*β* repertoires over time in healthy individuals we considered data from Chu et al. (*11*) which consists of TCR*β* sequencing data from serial blood samples collected at baseline and 1, 2, 3, 5, 6, 7, and 12 months after, from three healthy female volunteers (Figure 1A, dashed lines). Repertoire data was generated using the immunoSEQ assay (Adaptive Biotechnologies) with DNA derived from ∼10^6^ peripheral blood mononuclear cells (PBMC) (*11*). Plotting the frequency of clonotypes over time (“trajectories”) reveals a rich range of dynamic behaviour (Figure 1A). While some trajectories remain stable through time (grey data points), others exhibit large fluctuations, changing by up to 4 orders of magnitude over month-long timescales (pink and red data points). However, without a quantitative understanding of technical noise from repertoire sequencing, it is not clear to what extent these fluctuations are explained by statistical variance versus immune selection.

**Fig. 1.**
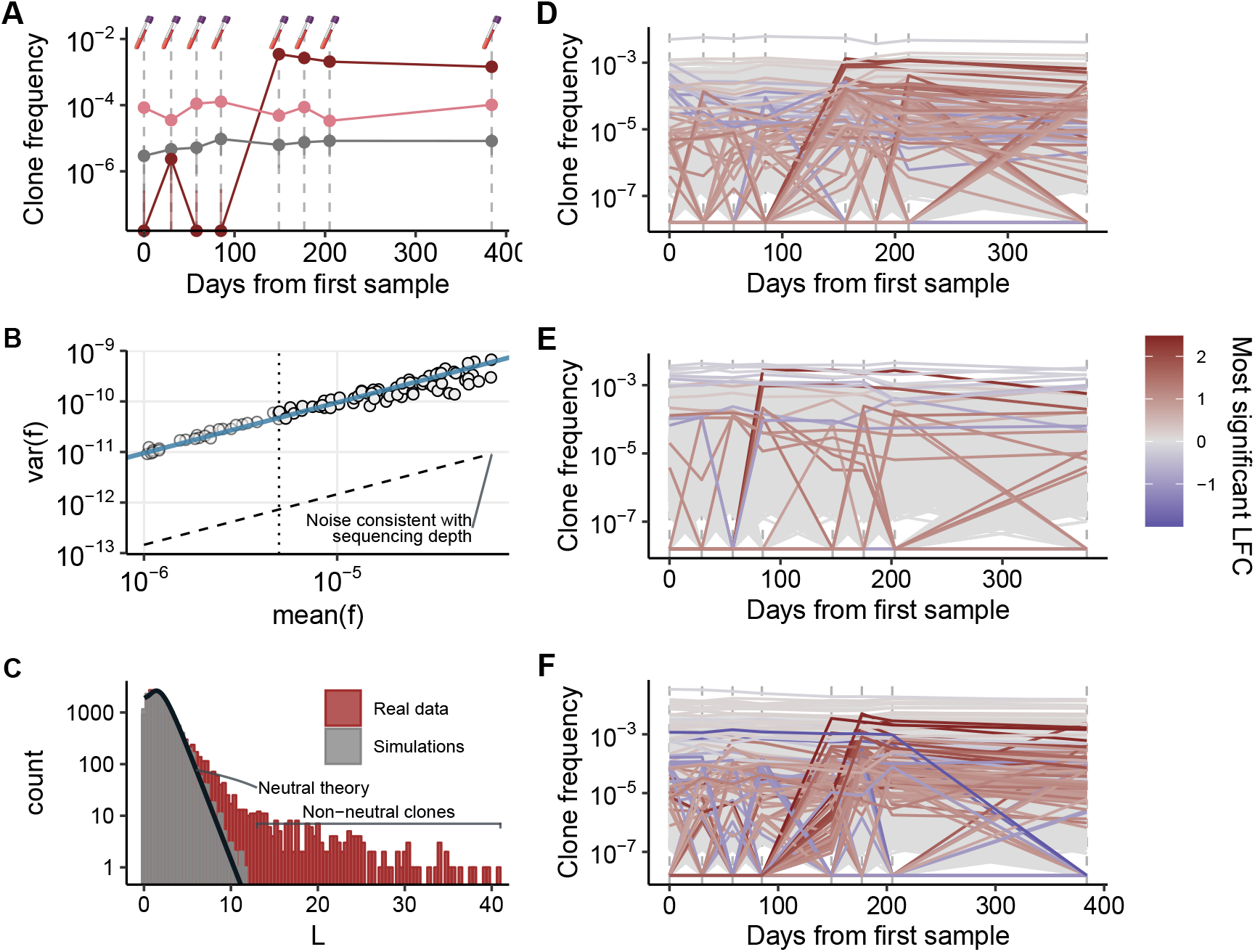
Identifying clonotype trajectories under immune selection. **A**. Example trajectories from Subject03 showing a clonotype exhibiting stable (grey), modest (pink), and large (red) temporal fluctuations in frequency. **B**. Variance in clonotype frequency across one pair of samples collected within 1 month of each other versus mean frequency shows the linear relationship (data points and best fit line) consistent with a sample size *N*_*e*_ substantially smaller than sequencing depth (dashed line). Linear fit excludes data points with very low frequencies (left of the dotted line) to remove discrete number effects. **C**. Distribution of *L* for simulated neutral trajectories (gray) and real data (red) across all values of *k*. Black solid line indicates the expected theoretical distribution. **D-F**. Clonotype trajectories across the 3 subjects where coloured trajectories are those with strong evidence of being under immune selection (FDR<0.1%). Colour indicates log-fold change (LFC) of the most extreme jump.

The experimental process of obtaining frequencies for each TCR clonotype effectively involves two sampling steps: one due to PCR and the other due to sequencing. A sample of blood containing 10^6^ PBMCs is expected to contain between 450,000 – 700,000 rearranged TCR*β* templates (*33*), a subset of which will be successfully amplified during multiplex PCR. Estimating clonotype frequencies from this amplified DNA using read counts introduces further sampling noise due to finite read depth. To determine the relative contributions from each of these two sampling processes, we carefully quantified levels of technical noise from samples collected within 1 month of each other (Figure 1B, Methods). Variance in clonotype frequency due to sampling noise is expected to scale linearly with frequency with an amplitude of ∼ 1*/N*_*e*_ where *N*_*e*_ is the effective sample size, resulting from the combination of both sampling steps. The variance observed from samples collected within 1 month of each other does indeed increase linearly with clonotype frequency, with effective sample sizes 100, 000 *< N*_*e*_ *<* 300, 000 – two orders of magnitude smaller than the sequencing coverage for this data. The fluctuations in frequency observed between samples collected within 1 month of each other closely matched the fluctuations observed in technical replicates (Supplementary Figure S1). This indicates that levels of technical noise are dominated by the finite number of rearranged TCR*β* templates captured during the multiplex PCR.

Having established that technical noise is dominated by fluctuations consistent with an effective sample size 100, 000 *< N*_*e*_ *<* 300, 000, we are able to develop a method to rationally identify trajectories that fluctuate by more than is expected due to sampling effects. We first downscale the sequencing data to *N*_*e*_ to ensure that variation in read counts dominates the technical noise (Methods). This enabled us to test how extreme a clonotype’s change in frequency is from one time point to the next by applying Fisher’s Exact Test to the down-scaled read counts. This down-scaling is crucial to avoid false positives resulting from assuming that the noise is driven by sequencing depth.

In order to integrate information about fluctuations in frequency across the entire longitudinal time-course, we considered the statistics of the most extreme jump in frequency along its trajectory. To do this we considered a statistic, *L*, which is the maximum of the − ln p-values, across each of the *k* recorded Fisher’s Exact Test p-values for a given clonotype:

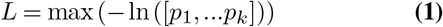

Large positive values of *L* indicate trajectories that have extreme jumps in frequency. To determine what value of *L* likely identifies trajectories whose fluctuations are unlikely to result from technical noise we developed an analytical expression for the distribution of *L* expected under neutrality (Methods). To check that this analytical expression does indeed capture the distribution of *L* expected under the null model, we simulated the trajectories of clonotypes across 8 time points in the absence of any immune selection (Methods). To closely match the real data, we sampled 100,000 – 300,000 clonotypes (modelling PCR) from an underlying frequency distribution that closely matches the observed distribution (Supplementary Figure S2) and then generated read counts for these clonotypes by subsequently sampling to a total depth of 10 million “reads” (modelling sequencing). These sequencing counts were then down-scaled following the same procedure as used for the real data. The observed distribution of *L* for these simulated data (Figure 1C, gray histogram, Supplementary Figure S3) closely matches our analytical expression (Figure 1C, black line). However, when the same procedure is performed on the real data, we observe a clear tail of non-neutral clonotypes (Figure 1C, red histogram, Supplementary Figure S3).

### TCR repertoires are largely stable over 1 year

Applying our framework to each subject we identified 30-140 clonotypes per subject whose trajectories showed evidence of immune selection over a period of one year (FDR=0.1%, Figure 1D-F). Across each subject a total of 6,234, 5,235 and 8,157 clonotypes passed our filtering (Methods) meaning that *>* 98% of clonotypes exhibit no evidence of changing frequency over the course of a year at this level of statistical power. The 0.5-1.7% of clonotypes showing evidence of being under immune selection sometimes show dramatic changes in frequency (up to 4-orders of magnitude over a 1-2 month time interval) and appear to change in coordinated waves of expansion or contraction (Figure 1D-F).

### Dynamic similarity and clonotype clusters

We reasoned that T-cells responding to a common stimulus might show highly coordinated dynamic behaviour. To assess how dynamically similar two clonotypes are through time we calculated the Pearson’s correlation coefficient, *r*, between their trajectories (Methods) and assigned a dynamic distance, *d*, between the trajectories using 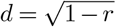. Trajectories exhibiting highly correlated dynamic behaviour have small dynamic distance (e.g. Figure 2A) while dissimilar trajectories have a large distance (e.g. Figure 2B and C). Using this distance metric we were able to build trees detailing the dynamic similarities of all sequences under immune selection in a subject (Figure 2D, Supplementary Figure S4). We observe a large fraction of clonotypes which coalesce at low branch heights, indicating a number of distinct clusters with highly similar dynamic behaviour. To determine how robust these dynamic clusters are, we considered an ensemble of 1,000 trees built by randomly permuting the time series for each clonotype (Figure 2E and F). Permuted trees show a much smaller fraction of clonotypes coalescing at low branch heights (Figure 2G) indicating that the strong groupings observed in the real data are unlikely to be a result of clustering noise. To determine a tree height threshold for identifying dynamic clusters we considered the height which maximised the difference in the fraction of clonotypes assigned to a cluster between the observed and permuted data (Figure 2G and Supplementary Figure S5).

**Fig. 2.**
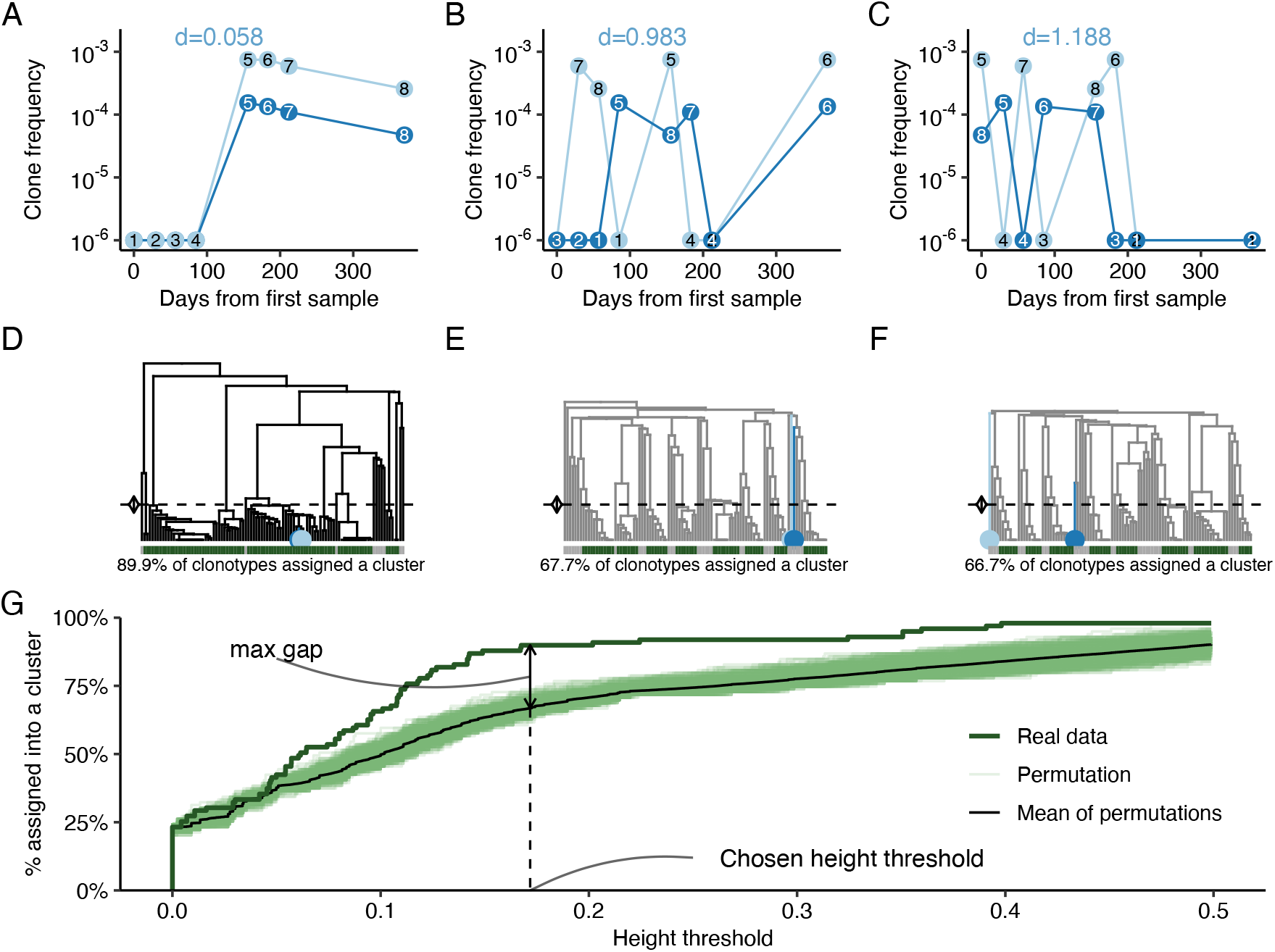
Identification of distinct dynamic clusters. **A**. Example trajectories of two correlated clonotypes, with a low dynamic distance *d*. **B-C**. Randomly permuted trajectories of the same two clonotypes, where the correlation is lost and the dynamic distance *d* is higher. **D**. Dendrogram built from all trajectories from clonotypes identified to be under immune selection in Subject01. Clonotypes in A. are highlighted with blue dots. Dashed line represents the chosen height threshold for clustering, and colours in the bottom bar show whether each clonotype has been assigned a cluster (green) or not (gray). **E-F**. Same as D., but corresponding to random permutations of each clonotype. **G**. The fraction of clonotypes assigned to a cluster as a function of tree cutting height threshold. The real data is a strong outlier relative to permuted data. The chosen height threshold (point of maximum difference) is shown by dashed line.

Using this approach we identified a total of 34 dynamic clusters across the three subjects (Figure 3 and Supplementary Figure S4). The dynamic clusters we identify are composed of between 2-36 clonotypes with the majority of rearrangements within a cluster being productive (Figure 3). In 11 out of the 34 dynamic clusters we observe a characteristic dynamics consisting of a rapid expansion in clonotype frequency (increasing by factors of between 10 and 1000-fold over <1-2 months) followed by a slower decay (decreasing by ∼ 10-fold over 4-8 months).

**Fig. 3.**
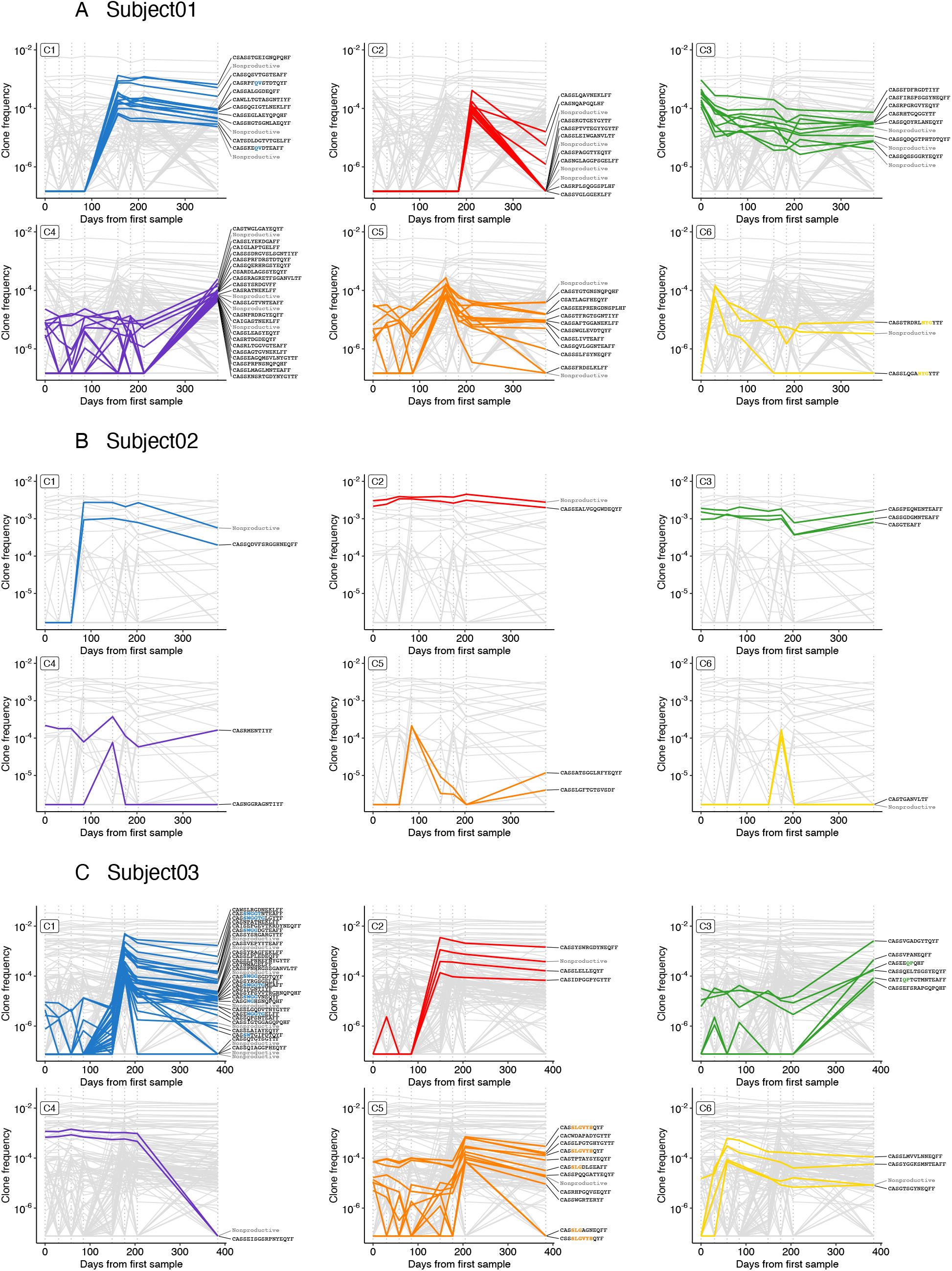
Clonotypes exhibiting highly similar dynamic behaviour show shared sequence features. (A-C) The top 6 most significant dynamics clusters (highlighted trajectories) across subjects 01, 02 and 03 respectively against the background of all sequences under immune selection in that subject (grey trajectories). CDR3 amino acid sequences are annotated on the right-hand side and statistically significantly enriched motifs (GLIPH2) are highlighted in bold colour.

### Does dynamic similarity imply sequence similarity?

We next considered whether there was evidence of sequence-level similarity for clonotypes exhibiting similar temporal behaviour (i.e. sequences within the same dynamic cluster). Previous work has demonstrated that shared biological function can manifest in enrichment for certain specific sequence motifs, especially in the complementarity-determining region 3 (CDR3) (*20, 29, 30, 34, 35*). We therefore used GLIPH2 (*29, 30*) to look for statistical enrichment of short amino acid motifs within dynamic clusters. We find that 6 out of the 34 dynamic clusters do indeed contain at least one enriched amino acid motif (*p <* 0.001, Fisher’s Exact Test), although these motifs are only seen in a subset of clonotypes within a cluster (Figure 3, Supplementary Figures S6, S7 and S8). Consistent with these motifs being a result of convergent biological function, the position of enriched motifs in the CDR3 sequence appears to be conserved across most clusters (Figure 3, Supplementary Supplementary Figures S6, S7 and S8) although more data is required to establish this conclusively. We hypothesise that the CDR3 sequences with the same enriched amino acid motif at similar positions within dynamic clusters are candidates for clonotypes which are recognising the same MHC-peptide complex.

### Does sequence similarity imply temporal similarity?

Next we sought to determine whether sequences with statistically significant enrichment of motifs showed evidence of shared dynamics. We reasoned that if the sharing of highly improbable motifs implies a shared antigen specificity, sequences with shared motifs would also show highly correlated dynamic behaviour. To test this we applied GLIPH2 to the CDR3 sequences from all clonotypes showing evidence of immune selection across each of the 3 individuals. This sequence-only based approach for identifying sequences under immune selection identified a total of only 24 sequences with statistically significant enrichment of 6 different motifs (*p <* 0.001, Fisher’s Exact Test). Surprisingly, however, we find that in only 2 of these 6 groups do a subset of the sequences belong to the same dynamic cluster (Figure 4 clusters A and C). In the other 4 groups the sequences show little evidence of shared dynamics: groups are composed of sequences that derive from different dynamic clusters (Supplementary Figure S4). This result highlights that, in some cases, sequences which share statistically significant motif usage may not in fact share antigen specificity as their dynamics show little evidence of being correlated.

**Fig. 4.**
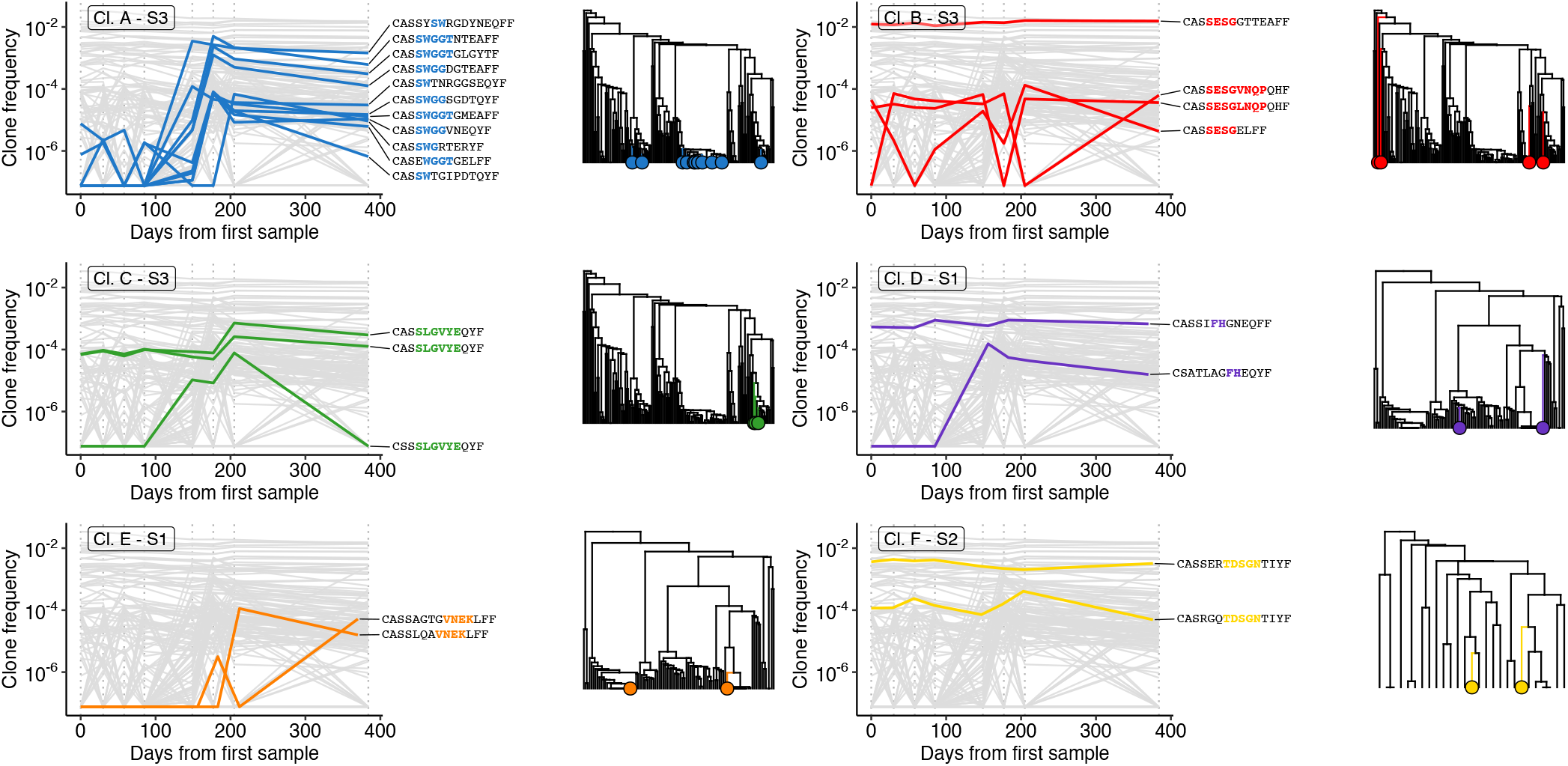
Non-neutral clonotypes sharing sequence motifs only partially show similar temporal behaviour. Sequence-based clusters identified by GLIPH2 on all non-neutral TCR sequences show a mixture of temporally coordinated and uncoordinated clonotypes.

### Hitchhiking of non-productive rearrangements

Five of the dynamic clusters identified consist of a productive CDR3 paired with a non-productive CDR3 (C8 in subject 1, C1, C2, C6 and C9 in subject 2 and C4 in subject 3). We reasoned that these pairs are likely to be examples of lineages in which V(D)J recombination resulted in a non-productive rearrangement with the cell subsequently producing a productive rearrangement by recombining the homologous chromosome (*2*). These pairs provide useful insights into systematic errors in TCR*β* frequency measurements because it is likely that the underlying frequency of both the productive and non-productive CDR3 sequence in the pair is the same. Despite TCR*β* sequencing approaches using spike-ins of synthetic templates with known frequencies in order to reduce biases (*36*), pairs of non-productive and productive CDR3s can differ in their frequencies by a substantial factor (e.g. C1 in subject 2), suggesting that considerable biases may remain likely resulting from differences in primer binding efficiencies in the early rounds of PCR.

## Discussion

Here we have developed a framework for identifying individual TCR clonotypes that are under immune selection in healthy individuals from longitudinal repertoire data. The basis of our method is to quantify levels of technical noise expected in the absence of immune selection and use this to identify clonotypes whose fluctuations exceed those expected due to statistical noise. By applying this framework to TCR repertoires from three healthy individuals collected over a period of a year across 8 serial samples, we show that the majority of the repertoire is highly stable. However, a minority of clonotypes exhibit highly coordinated waves of immune selection and in some of these waves, there is evidence of shared antigen specificity.

Because the TCR repertoire is so diverse, the majority of clonotypes exist at low frequencies and are sampled in modest numbers of sequencing reads. In data such as these, it is crucial to accurately understand the fluctuations expected due to technical noise. We have shown that sequencing read count data can underestimate the expected fluctuations in clonotype frequencies. By carefully considering the frequency fluctuations observed across neighbouring time points and technical replicates, we have shown that for these data fluctuations are driven by the number of template molecules captured in the initial stages of PCR rather than by sequencing depth. To rationally detect clonotypes that are systematically expanding or contracting, estimates of the number of templates captured are needed rather than raw sequencing coverage. For most data this also implies that more accurate estimates of clonotype frequencies require larger quantities of DNA at the outset of PCR rather than deeper sequencing. A similar framework developed for RNA based RepSeq data (*32*) recently inferred that technical noise is consistent with template copies of between 10^6^ − 10^7^. This may indicate that the number of templates captured in RNA based RepSeq data could be substantially higher than for DNA thus reducing technical noise. However, it is not clear that these more accurately reflect true clonotype frequencies as expression levels may vary substantially from clonotype to clonotype.

The clonotypes that our framework identifies as being under immune selection fall into clear dynamic clusters with clonotypes showing highly similar temporal patterns. These dynamic patterns often have a characteristic expansion-decay profile which is captured over the course of a year (e.g. C1 subject 3). Accurate estimation of the expansion / decay kinetics is challenging because of the non-uniform time intervals between samples and because of it is often not possible to observe the clonotypes at their pre-expansion levels due to finite depth (e.g. C2 subject 1). However it appears that expansion is more rapid than decay. We find that the number of unique TCRs in a dynamic cluster is smaller than the numbers reported from vaccine studies (*19*). This may be because the unknown stimuli are less immunogenic than vaccines, or because many more clonotypes exhibiting similar dynamic behaviour exist at frequencies below our detection limits.

Previous work has shown that certain CDR3 sequence motifs have been linked to shared functional specificity of the TCR (*29, 30*). We find a handful of examples where shared dynamics also manifests in shared similarity at the sequence level with conserved motifs at similar positions in the CDR3 sequence. However we also find many examples of sequences exhibiting highly similar dynamics with no apparent sequence level similarity. This observation is consistent with the fact that shared motifs will only be observed in TCRs that bind the same antigen, however a given stimulus (e.g. a virus) likely presents many different antigens that are recognised by TCRs. Indeed with improved data it may be possible to estimate how many distinct antigens are eliciting an immune response by looking for distinct clusters of CDR3 sequences within the same dynamic cluster.

In some cases, pairs of sequences exhibiting highly similar dynamic behaviour are composed of a non-productive sequence paired to a productive one. Previous single-cell studies have found that a second V(D)J recombination can happen in a T-cell after the first resulted in a non-productive rearrangement, and that this is not a rare occurrence (*2*). The existence of such pairs with highly similar dynamics observed in the data here strongly suggests they are in the same clone and would thus have the exact same underlying frequency. However, the estimated frequencies in these pairs can differ by an order of magnitude. This provides evidence that, despite the use of synthetic templates as an error correction strategy to reduce systematic biases (*36*), it is likely some systematic biases remain in the frequencies reported from the Adaptive Biotechnologies platform.

Better characterisation of immune repertoire dynamics in healthy individuals could be achieved using blood samples collected with high temporal resolution (e.g. each month) over many years from large numbers of healthy individuals. For example, 10mL of peripheral blood could provide a sample of ∼ 10^7^ T-cells which would provide a resolution two orders of magnitude better than in the data reported here. This deeper characterisation of the repertoire (including potentially linking some of the TCR*α* and TCR*β* chains) could then provide a more quantitative understanding of immune repertoire dynamics in the absence of overt disease. This characterisation of the “healthy background” dynamics, combined with a better understanding of which antigens CDR3 sequences are recognising, could have potential as a powerful detector of certain diseases.

## Methods

### Data

We analyse data generated by Chu et al. (*11*). The dataset comprises of immunosequencing of DNA extracted from peripheral blood mononuclear cells (PBMC) of 3 healthy individuals (all 3 of them women, aged 18-45). For each individual, blood was taken at 8 time points in the span of 12 months (at starting date and after 1, 2, 3, 5, 6, 7, and 12 months). The dataset consist of read count matrices for all unique nucleotide level CDR3 rearrangements sequenced, along with information about the V, D and J genes used, when known. The frequency of each rearrangement is calculated by dividing the correpsonding read count by the total read count for the sample. All samples were sequenced at a depth of 1 − 3 × 10^7^ total reads. Technical replicate data was obtained from Rytlewski et al (*10*).

### Filtering out contamination between samples

While checking for the typical overlap between intra- and inter-individual TCR repertoires, we found some evidence of contamination between some of the samples, as shown in Supplementary Figure S9. We observed that the last time point of subject 01 showed a substantial overlap with all repertoires from subject 03, and in particular time point 6 (8th row, 3rd to last column in Supplementary Figure S9), indicating that a fraction (roughly 3-4%) of the reads in time point 8 from subject 01 comes from time point 6 in subject 03. To control for this, we decide to remove all reads from time point 8 in subject 01 that also appear at higher frequencies in time point 6 from subject 03.

We also observe smaller instances of potential contamination in several samples, which on the 2D heatmaps look like small clusters of points in the top left or bottom right corners of the plot, with some level of correlation in between samples. These features come from groups of shared rearrangements that exists at high frequency in one sample, and appear at low frequency in another sample, but with certain correlation on the frequency values. We filter these instances out by finding all rearrangements that appear in any pair of inter-individual samples where the frequency in one sample is at least 100× the frequency in the other one, and removing these rearrangements from the low frequency sample.

These filtering steps remove 31, 312 unique rearrangements out of the observed ∼ 9, 120, 000 across all 24 samples (0.34%).

### Quantifying technical noise and effective sample size

When working with read count data, binomial or Poisson approaches are commonly used, with a core assumption that the variance of the data will be consistent with a sample size equal to the sum of all reads (∼ 10^7^ reads in these data). We tested this assumption by exploring how the variance in frequency of a clonotype 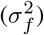 scales with its estimated frequency (*f*). For sampling noise the variance in frequency is expected to be linearly related to the frequency via 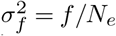 where *N*_*e*_ is the effective sample size driving noise. To check this relationship we bin clonotypes based on their frequency in one of the time points (or replicates) and calculate bin means. We can then estimate the variance in frequencies by considering the variance in the frequencies of these clonotypes in the other time point (or replicate). We can then estimate the effective sample size by using

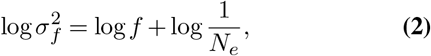

and plotting the log of the variance in frequency as a function of the log of the mean frequency as shown in Supplementary Figure S1. The effective sample size can then be estimated from the intercept of the best fit line on the plot. We performed this procedure first by considering technical replicates (Supplementary Figure S1) and second by considering sequential time points (main text Figure 1B).

An additional consideration we had to take when comparing different time points to estimate *N*_*e*_ is that we expect clonal expansion events over time whereby, for a given clone, the frequency will change drastically between samples which can cause some of the bins to have anomalously large variance (see outlier data points in Supplementary Figure S11). To minimise the effects of clonotype expansions artificially driving up estimates of the variance, we implement an extreme outlier filtering step. We first bin all clonotypes based on the average frequency across both samples 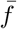. Then, for each bin of size *n*, we calculate an upper and lower threshold of clone frequency beyond which we would expect to observe zero clones from those bins, assuming the noise around 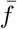 is consistent with a sample size of 200, 000. We remove any clonotype beyond these thresholds and then proceed to fit a linear model to estimate *N*_*e*_ as shown in Supplementary Figure S11.

### Statistical test for detection of non-neutral clones across the time series

For each comparison between subsequent time points, we use the calculated *N*_*e*_ as a downscaling factor for repertoire read counts. We obtain a corrected read count value by multiplying the frequency of each rearrangement by *N*_*e*_ and rounding it to the closest integer. This results in a dataset that is consistent with the assumptions for many count based statistical methods and ensures that the vast majority of the noise between samples is consistent with the downscaled read count and controls the false positive rate.

We then perform Fisher’s Exact Test for the read counts of all clonotypes present in at least one of the two samples, filtering out any clonotype with less than 6 read counts across both downscaled repertoires. We do this for all 7 pairwise comparisons between subsequent time points in each individual, resulting in *k* = 1, 2, …, 7 p-values per clonotype, depending on how many time points each clonotype is detected in.

To summarise the results of all *k p*-values corresponding to a clonotype into a single overall significance score, we calculate *L* = max (− ln ([*p*_1_, …*p*_*k*_])). This metric favours the detection of trajectories where a sudden clonal expansion taking place between two time points results in a very low *p*-value for that step, but it can also capture more subtle trajectories. Importantly, because *p*-values follow a standard uniform distribution under the null, − ln *p* will follow an exponential distribution with rate *λ* = 1. As a result, we can derive a theoretical expectation for *L* as the expected distribution of the maximum of *k* i.i.d. exponential distributions:

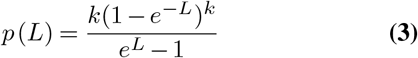

This null model allows us to obtain an aggregated *p*-value that summarises the entire trajectory, being able to detect non-neutral clones across the time series. We lastly correct for multiple testing using the Benjamini–Hochberg procedure to obtain a False Discovery Rate (FDR). We select for further analysis trajectories with a FDR*<* 0.1%.

### Simulations to assess false discovery rate

To evaluate the validity of our framework under the null, we simulated a time series of 8 TCR repertoires that show no changes in frequency over time. For each time point, we performed these simulations by first drawing 100, 000 – 300, 000 “rearrangements” from a reference set of rearrangements with distribution of true clone frequencies following a power law such as *p*(*f*) = 1*/f* ^2^. This step simulates the sampling of a finite number TCR rearrangements from the blood and their capture in the first PCR cycles during library preparation. The captured molecules are then “sequenced” to a depth of 10^7^ to obtain TCR repertoires and their read counts resembling real data.

We apply all the steps of our analysis framework to the simulated data: estimation of *N*_*e*_ in subsequent time points and corresponding downscaling of read counts, Fisher’s Exact Test and aggregation of resulting *p*-values to obtain *L*. By comparing the resulting distribution of *L* with its theoretical expectation under the null, we check that the model is valid (Supplementary Figure S3 and it does not lead to a higher False Positive Rate than expected (Supplementary Figure S10).

### Measuring dynamic similarity

For all the clonotypes detected as non-neutral at the chosen significance level in an individual, we calculated a distance matrix with a metric we define as dynamic distance 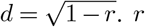 corresponds to Pearson’s correlation coefficient of the two vectors consisting of the raw frequencies at all 8 time points (assigning a frequency of 0 when a rearrangement is missing from a time point). Other similarity metrics are possible however we opted using one based on Pearson correlation as these have been previously used to detect similar trajectories in time series data e.g. Tikhonov et al. ISME (2015).

### Hierarchical clustering based on dynamic distance

From the obtained dynamic distance matrices we performed agglomerative hierarchical clustering, using single-linkage criteria. This resulted in one dendrogram per individual.

To rationally choose a height threshold for tree-cutting and obtaining distinct clusters, we performed a permutation based analysis, as shown in Figure 2. For each individual, we randomly permuted the trajectory of each clonotype, calculated the dynamic distance matrix and built a dendrogram by hierarchical clustering. Clusters derived from these dendrograms will not reflect real groups of clonotypes, but the background level of cluster formation for this type of data. By using the percentage of clonotypes in a subject assigned into a cluster as a metric, we studied the effect of varying tree cutting height thresholds on this metric. The chosen height threshold corresponds to the the height at which the difference between the real dendrogram and the mean of the permuted dendrograms is largest.

### Identification of sequence motif clonotype clusters

We applied GLIPH2 (*30*) to the set of productive CDR3 rearrangements from each dynamic cluster. GLIPH2 identifies for statistically entriched short amino acid motifs (2-5 amino acids) with respect to a reference set of CDR3 sequences, in our case the collection of all unique productive CDR3 rearrangements in each individual. Enrichment is determined by Fisher’s Exact Test, and we determine a threshold of *p <* 10^*−*3^.

## Code availability

All code used in this study is available on the Blundell laboratory GitHub page: https://github.com/blundelllab/TCR-dynamics.

## ACKNOWLEDGMENTS

We would like to thank Elizabeth Soilleux, Doug Easton, Daniel Fisher and members of the Blundell lab for helpful comment on the manuscipt.

## Data and code availability

All data and code used in this study will be made available on the [Blundell lab Github page](https://github.com/blundelllab/Clonalhaematopoiesis.git).

## Supplementary Figures

**Fig. S1.**
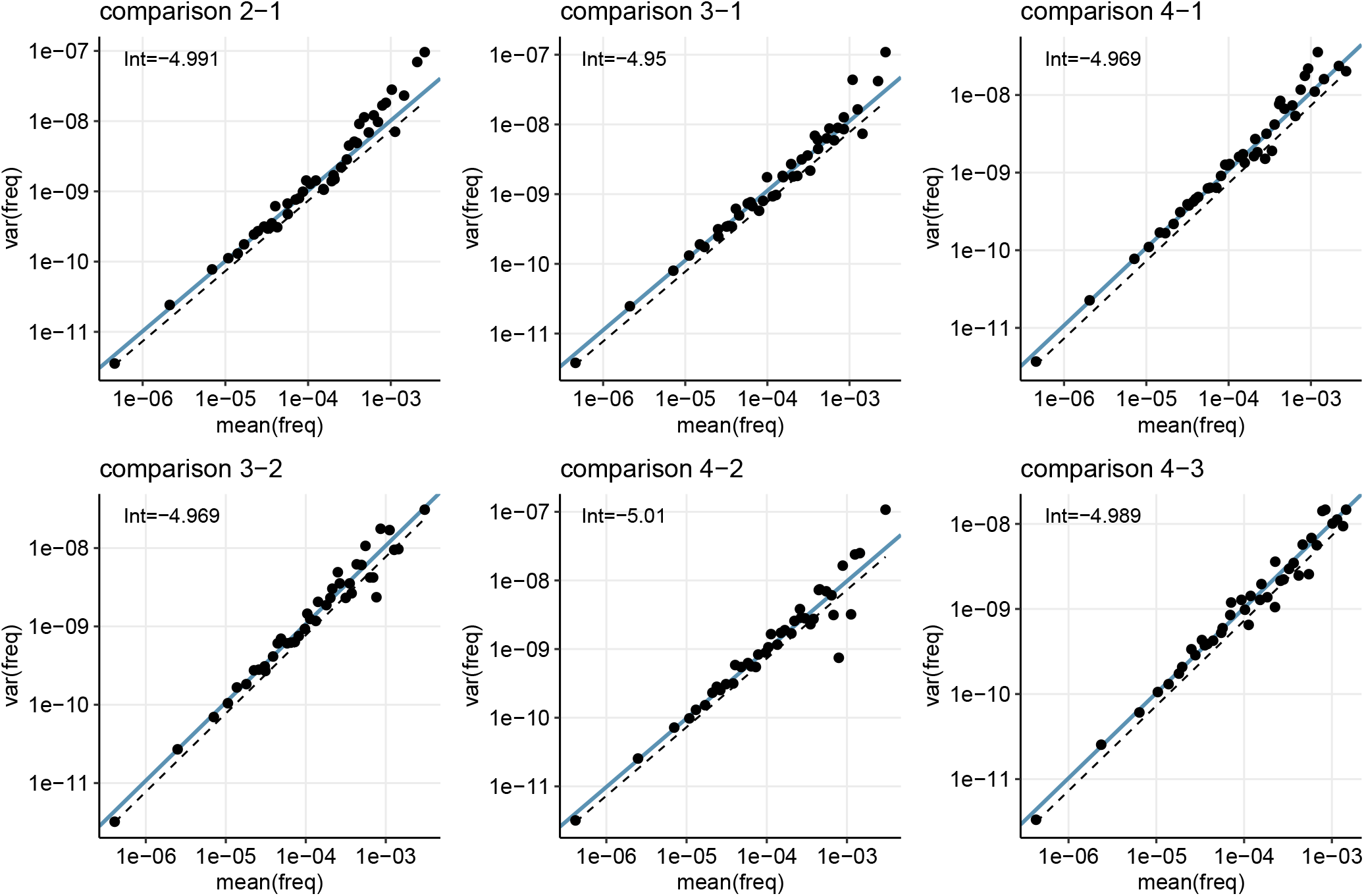
Technical replicates. Relationship between mean and variance in 4 technical replicates from Rytlewski et al. (*10*) (showing all possible pairwise comparisons) suggests an effective sample size of ∼ 200, 000 TCR molecules, as *N*_*e*_ = 2 × 10^*−*Intercept^

**Fig. S2.**
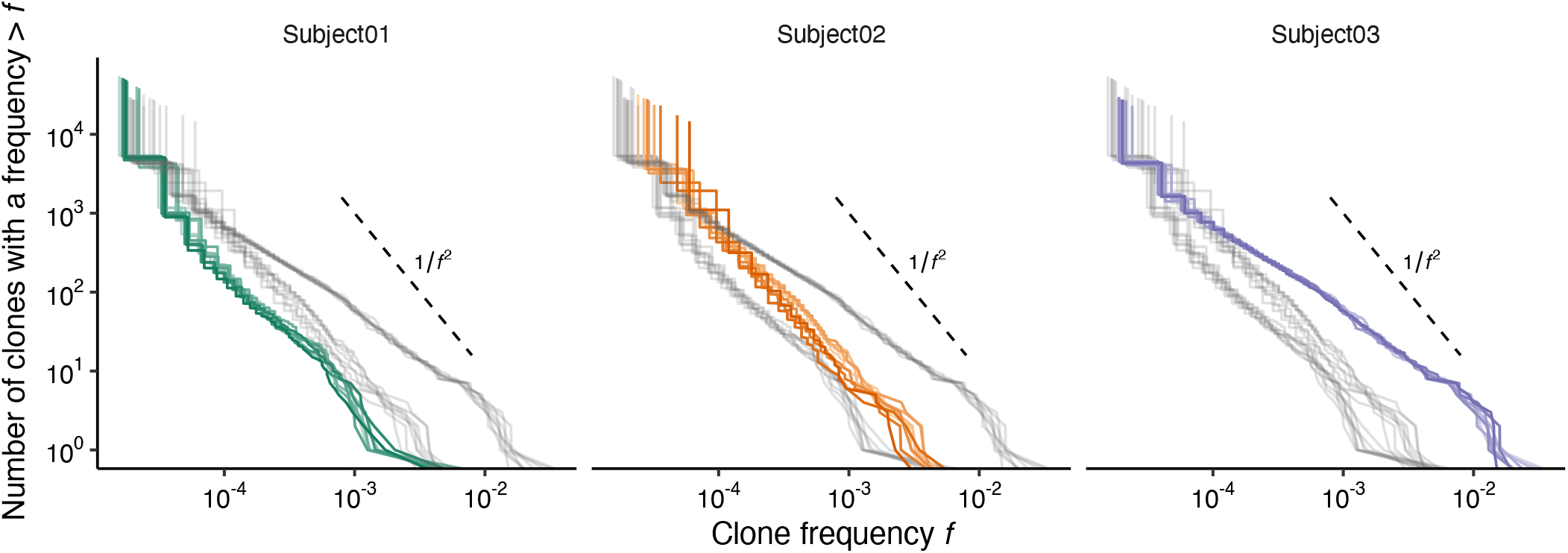
Distribution of clone sizes across all time points. Reverse cumulative plots for the clone frequency distribution in each time points (different shades of colour - lighter is earlier, darker is later) for all three individuals. All samples show a clear power law distribution, with almost no within-individual variation, but a clear difference between individuals. A reference is shown for the power law distribution *p*(*f*) = 1*/f* ^2^.

**Fig. S3.**
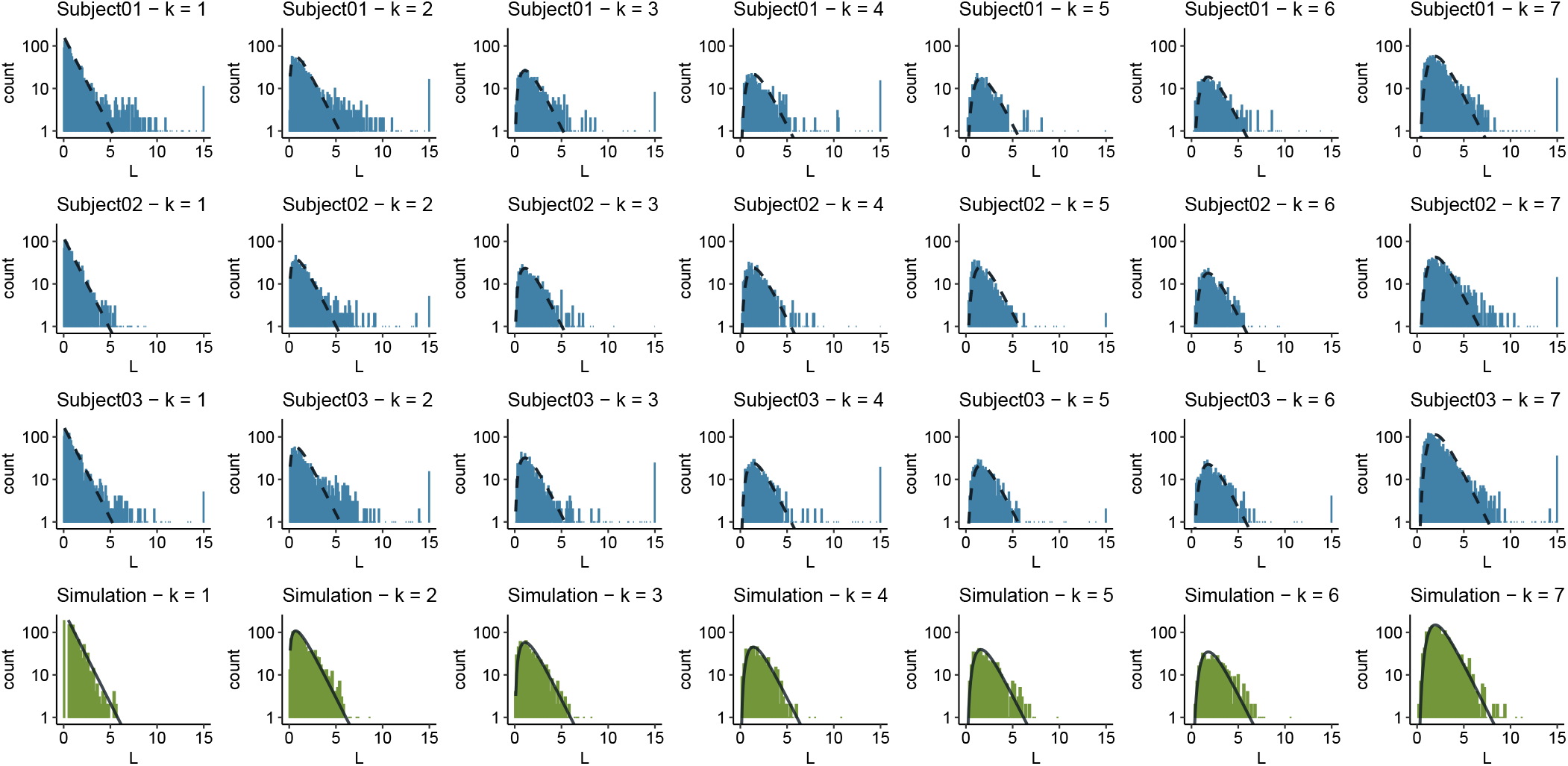
**Distribution of** *L* **for different numbers of available** *p***-values (***k*) in each one of the 3 individuals (blue) and simulations (green). Values beyond the upper limit of *L >* 15 have been condensed into the uppermost bin.

**Fig. S4.**
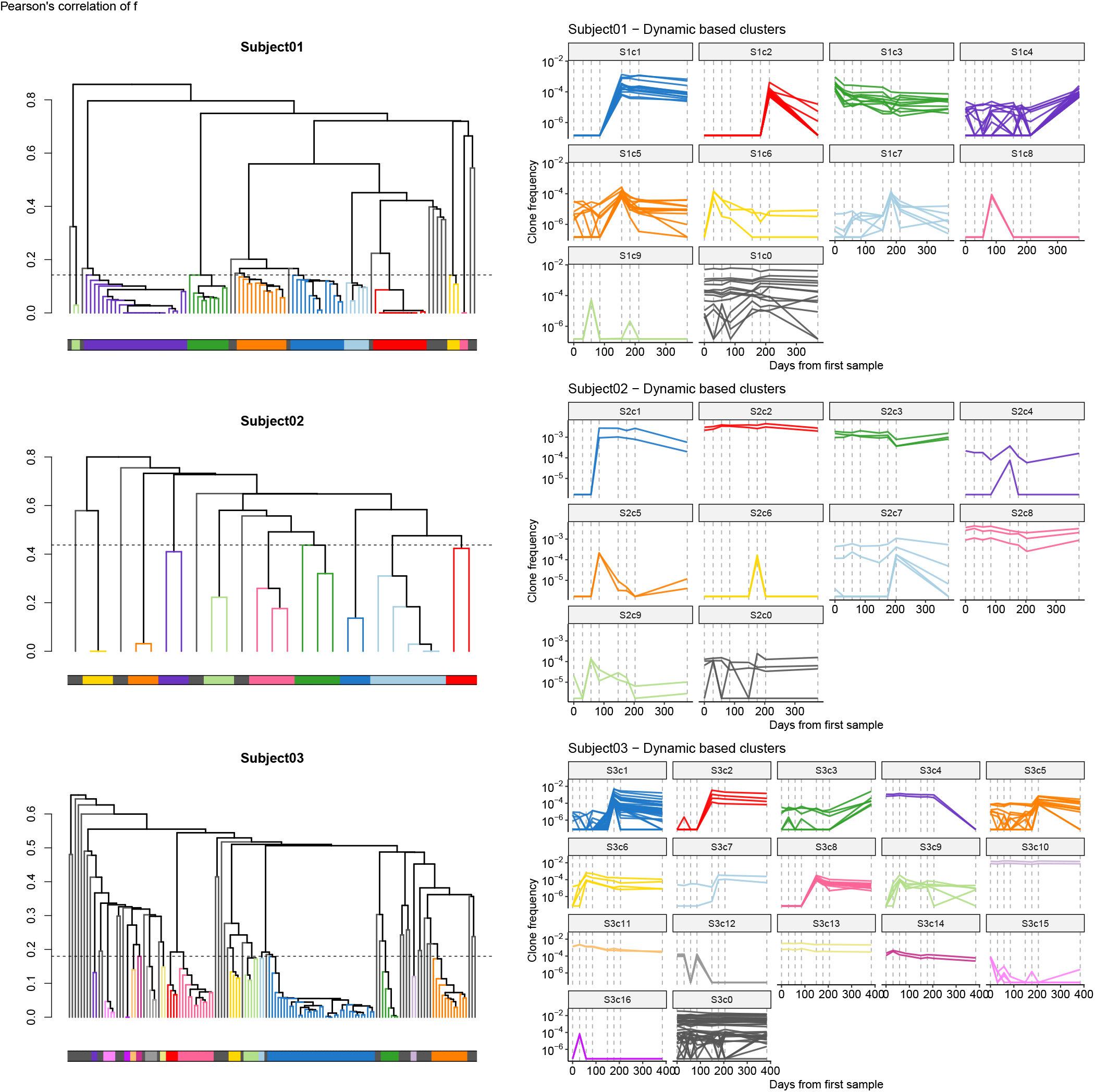
Hierarchical clustering based on dynamic distance identifies distinct dynamic clusters. **Left**. Dendrograms built from dynamic distance matrices from each individual, with the chosen height threshold (dashed line) for tree cutting and resulting clusters (colours). **Right**. Trajectories of the dynamic clusters, along with the trajectories of the outgroups (cluster 0, gray).

**Fig. S5.**
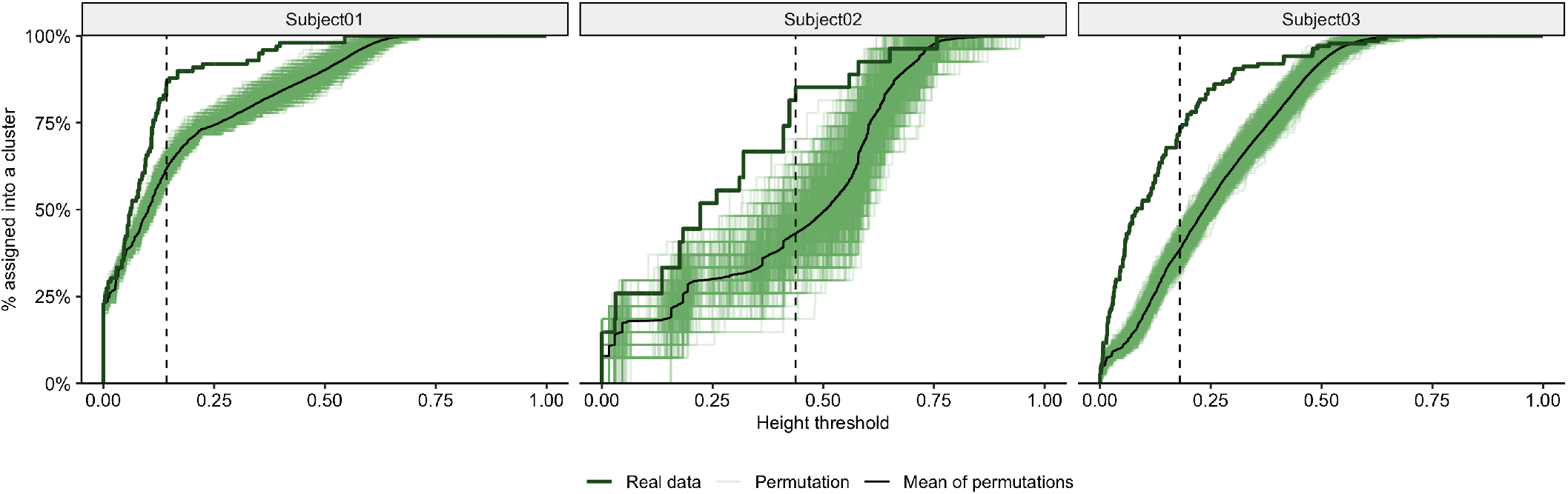
Finding a rational tree cutting height threshold. For the set of all non-neutral clonotypes in each individual, we performed 1,000 permutations of their trajectories and built dynamic distance trees. By considering different tree cutting height thresholds (x-axis), we studied the total fraction of clonotypes being assigned to a cluster (y-axis), and chose the height threshold at which the gap between the mean of the permutations (light green lines, average in black) and the real data (dark green line) is maximum.

**Fig. S6.**
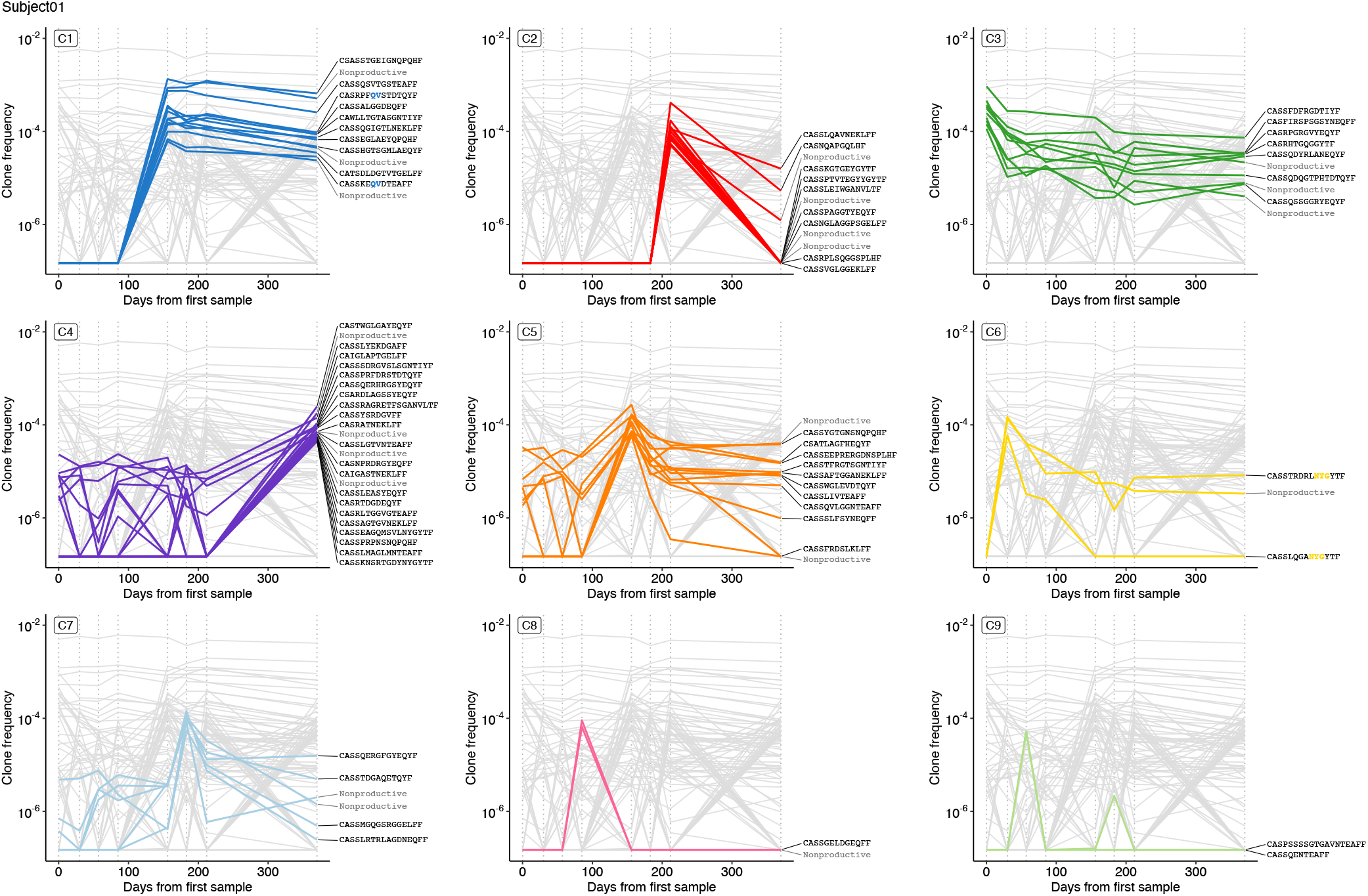
Dynamic clusters in Subject 01 along with their CDR3 amino acid sequence (if productive) and GLIPH2 identified sequence motifs highlighted in colour.

**Fig. S7.**
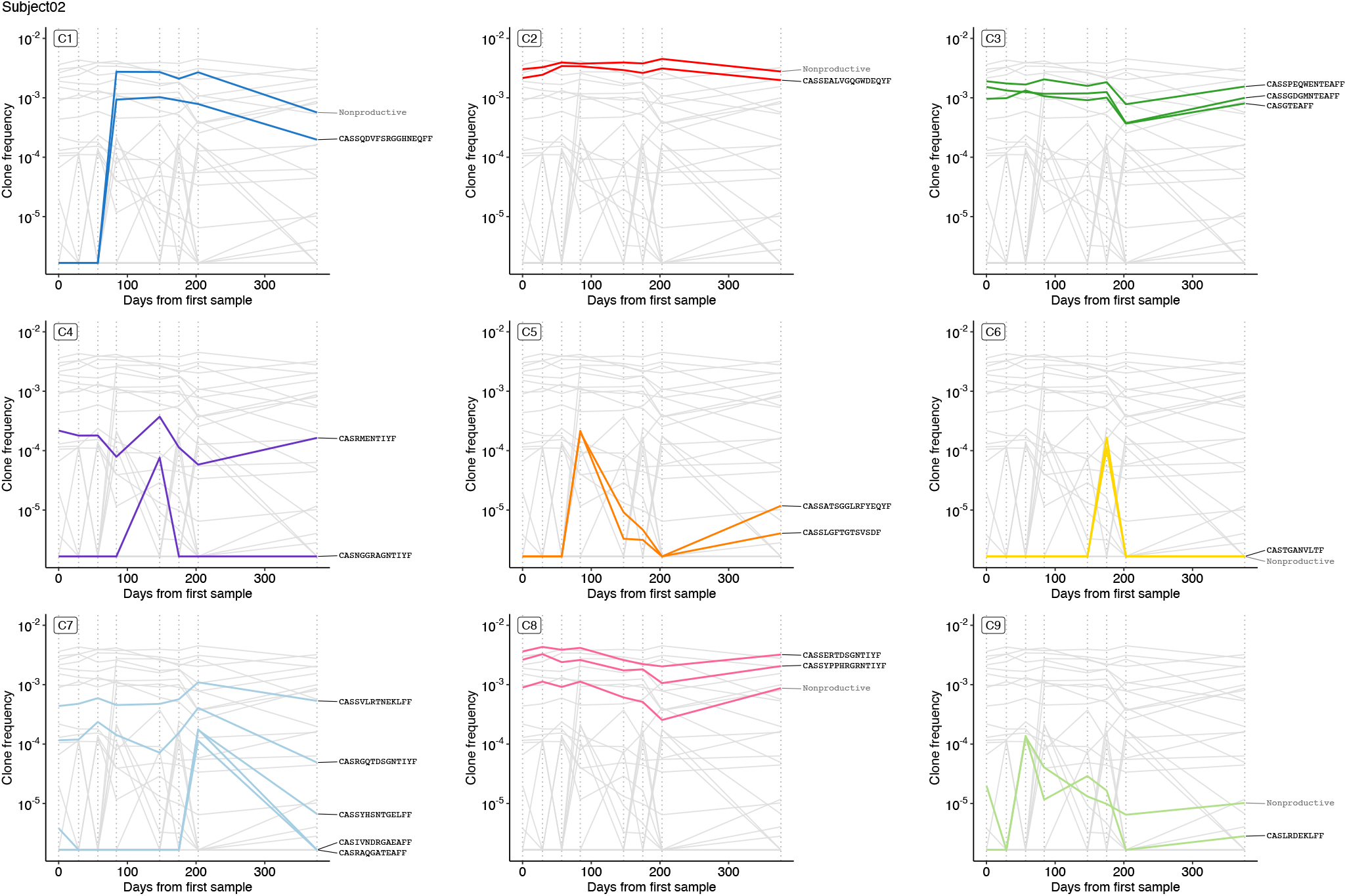
Dynamic clusters in Subject 02 along with their CDR3 amino acid sequence (if productive) and GLIPH2 identified sequence motifs highlighted in colour.

**Fig. S8.**
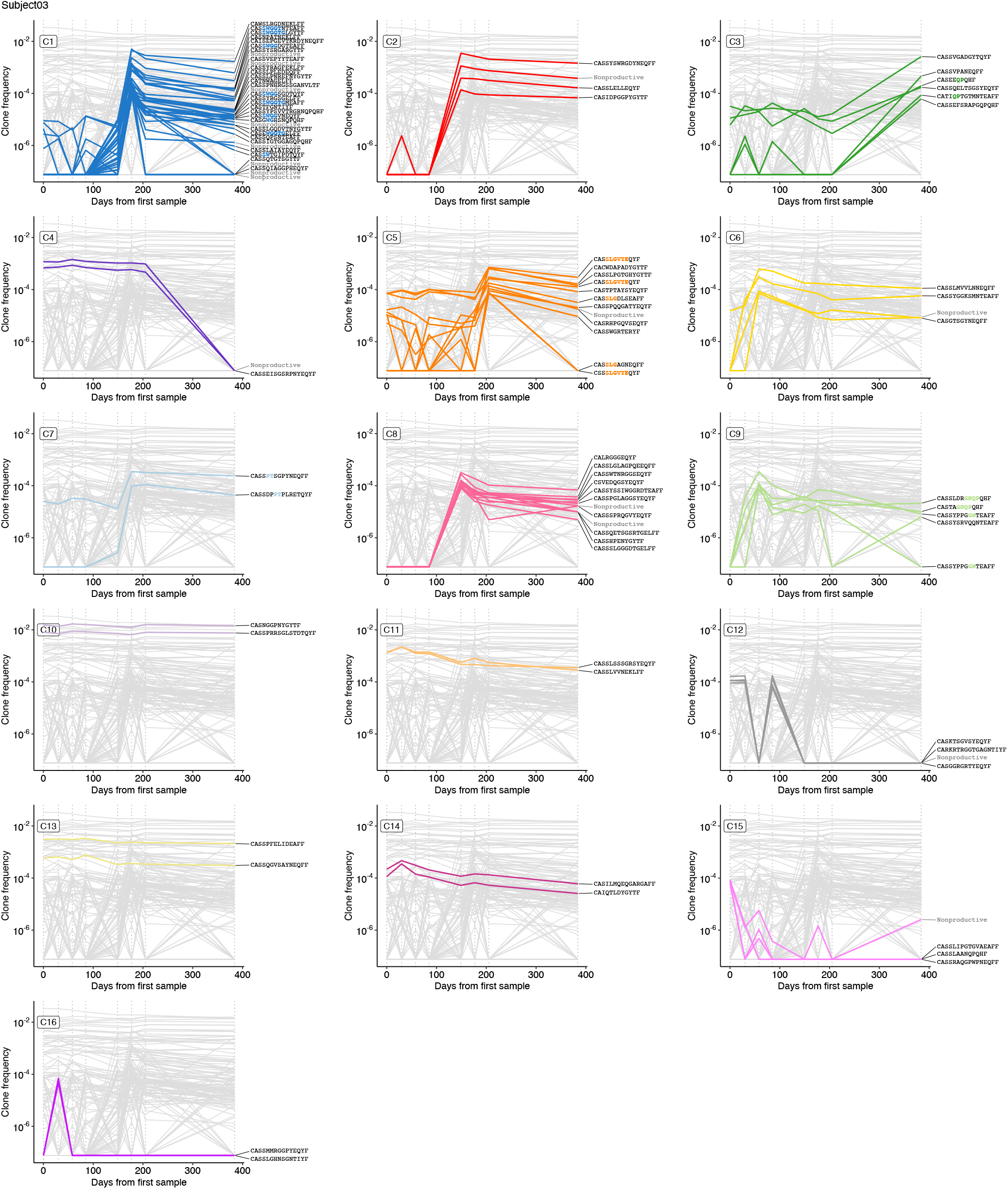
Dynamic clusters in Subject 03 along with their CDR3 amino acid sequence (if productive) and GLIPH2 identified sequence motifs highlighted in colour.

**Fig. S9.**
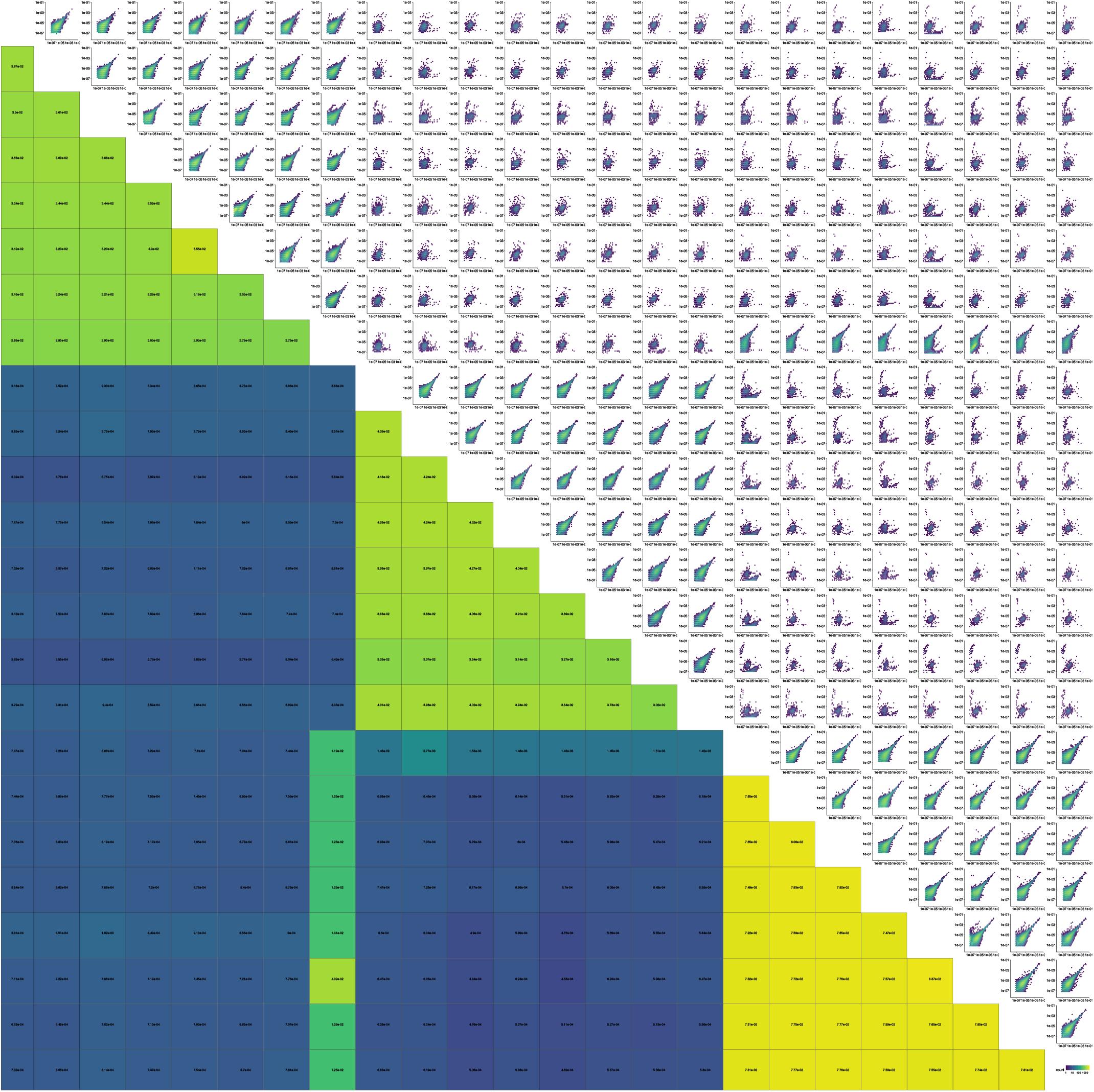
Evidence of contamination between repertoires. Rows and columns correspond to subjects 01, 02 and 03, each one in temporal order. **Upper triangle:** For every possible pair of samples, hexagonal 2D heatmap of the frequencies of overlapping TCR rearrangements on a nucleotide level. **Lower triangle:** Jaccard index (size of intersection over size of union) for each pair.

**Fig. S10.**
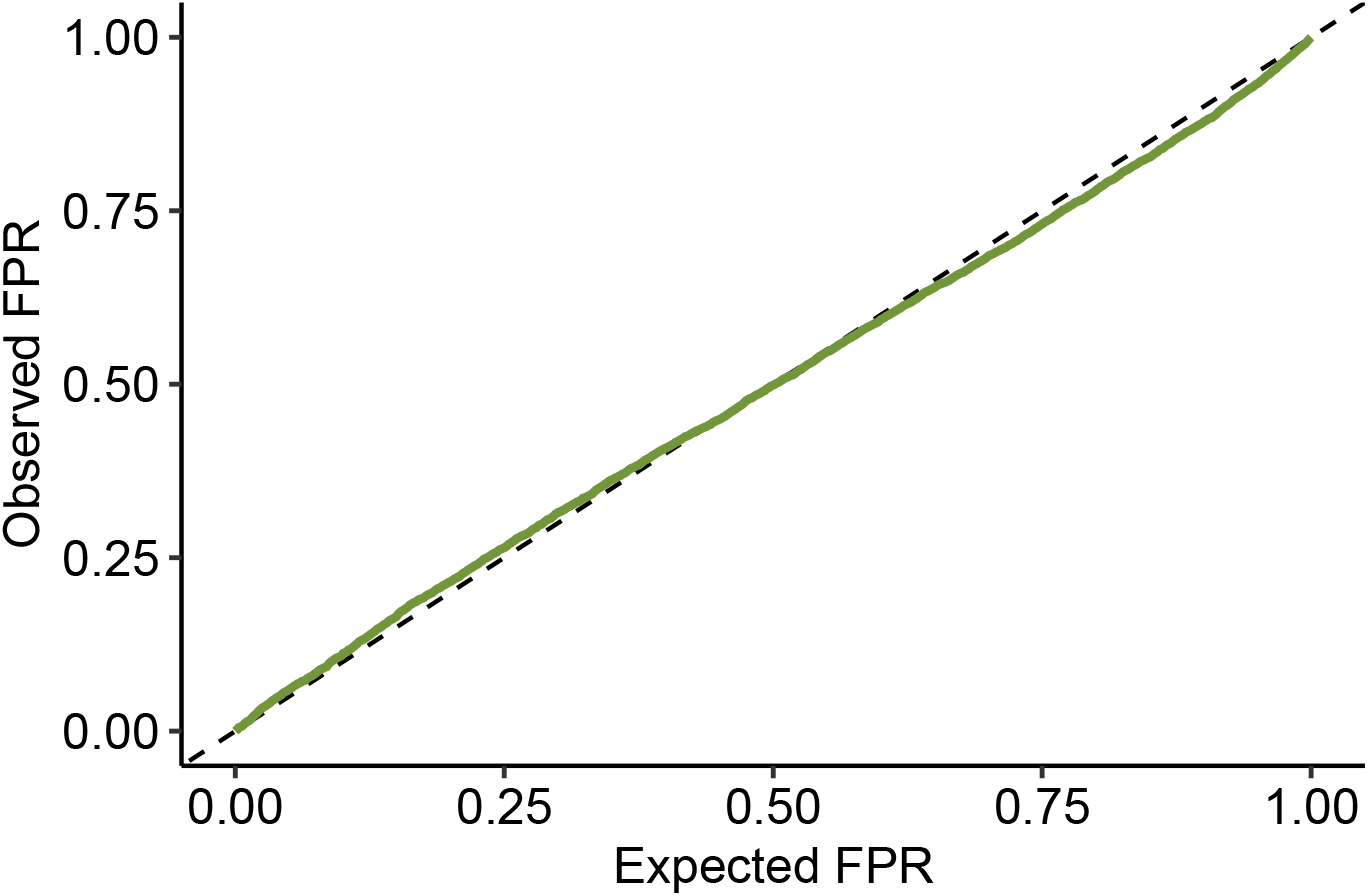
Expected False Positive Rate (FPR) matches the empirical FPR in null simulations. Using the null distribution of *L*, we can obtain a *p*-value per clonotype. For a given *p*-value threshold or *α* (Expected FPR), we count how many simulated clonotypes are identified as statistically significant (false positives) and divide the number by the total number of clonotypes, resulting in the Observed FPR.

**Fig. S11.**
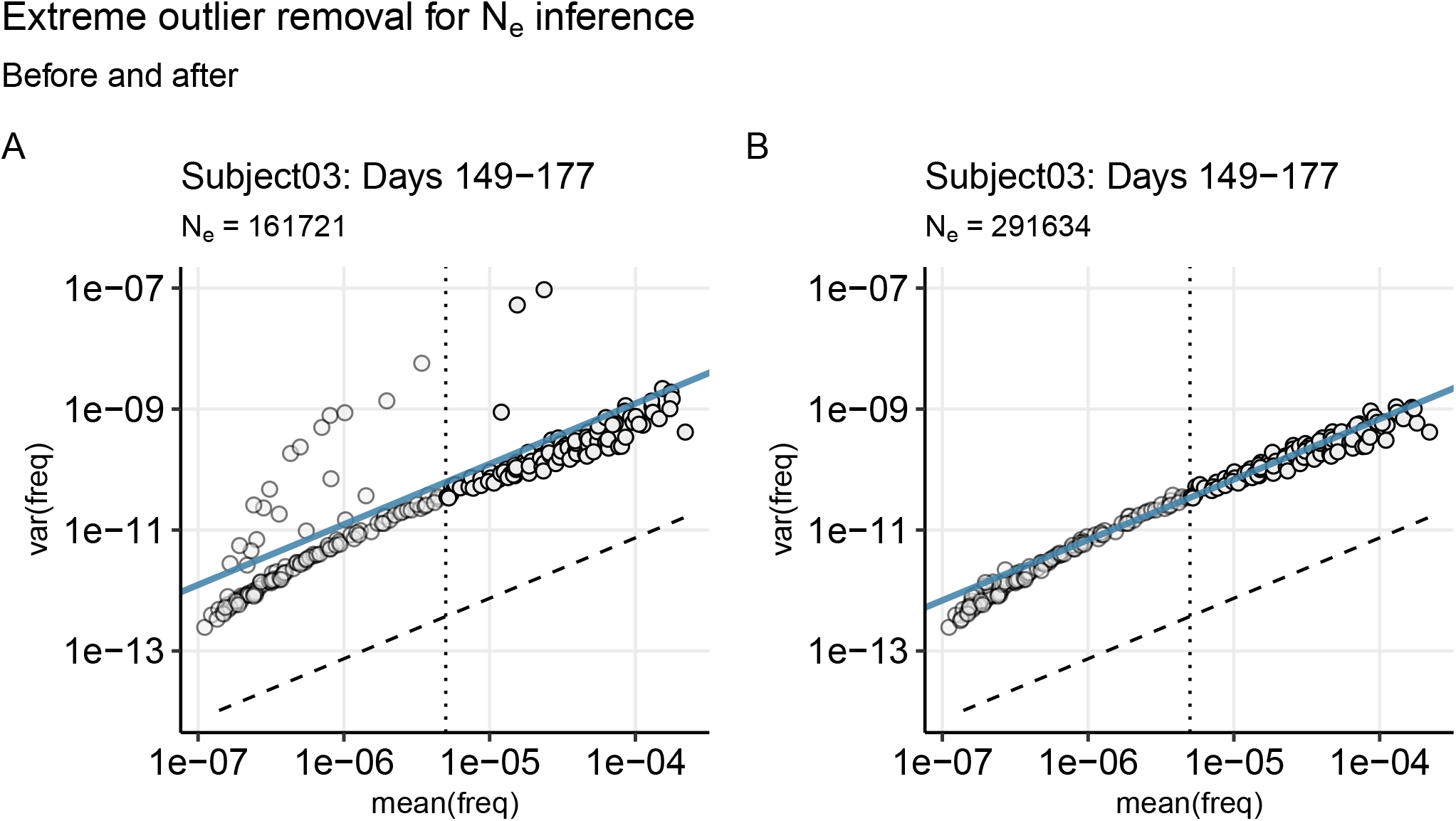
Effect of outlier removal on mean-variance scaling between two subsequent time points. Variance in clonotype frequency across one pair of samples collected within 1 month of each other versus mean frequency shows the linear relationship (data points and best fit line) consistent with a sample size Ne substantially smaller than sequencing depth (dashed line). Linear fit excludes data points with very low frequencies (left of the dotted line) to remove discrete number effects. **A**. Clonal expansions between one time point and the next will result in unusual outlier data points that overestimate the null variance for those bins. **B**. After filtering out extreme outliers, the linear fit is more unbiased.

